# Neural data science: accelerating the experiment-analysis-theory cycle in large-scale neuroscience

**DOI:** 10.1101/196949

**Authors:** L Paninski, J.P Cunningham

## Abstract

Modern large - scale multineuronal recording methodologies, including multielectrode arrays, calcium imaging, and optogenetic techniques, produce single - neuron resolution data of a magnitude and precision that were the realm of science fiction twenty years ago. The major bottlenecks in systems and circuit neuroscience no longer lie in simply collecting data from large neural populations, but also in *understanding* this data: developing novel scientific questions, with corresponding analysis techniques and experimental designs to fully harness these new capabilities and meaningfully interrogate these questions. Advances in methods for signal processing, network analysis, dimensionality reduction, and optimal control – developed in lockstep with advances in experimental neurotechnology - - promise major breakthroughs in multiple fundamental neuroscience problems. These trends are clear in a broad array of subfields of modern neuroscience; this review focuses on recent advances in methods for analyzing neural time - series data with single - neuronal precision.

**Figure 1.**
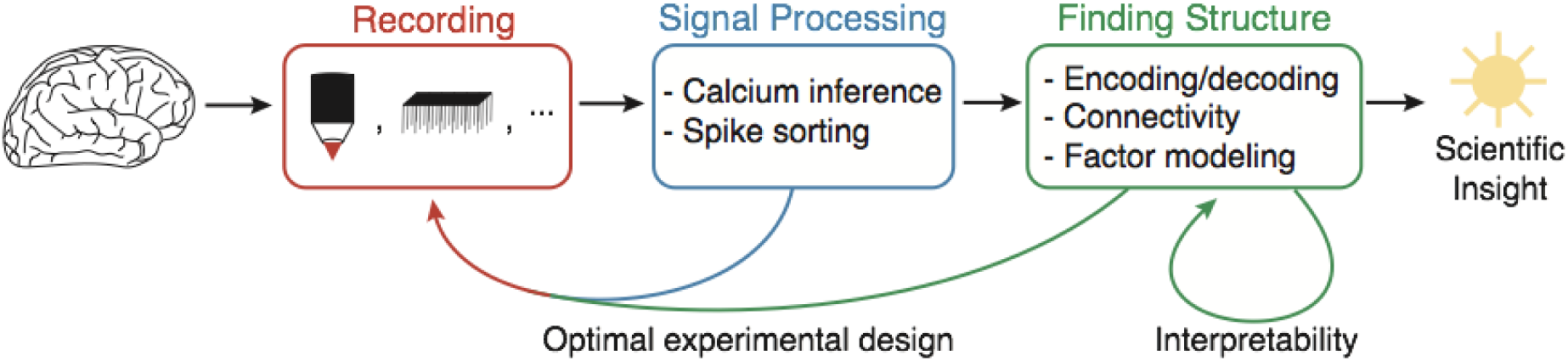
The central role of data science in modern large - scale neuroscience. Topics reviewed herein are indicated in black.

## I. High - throughput neural signal processing methods

Neuroscientists have long dreamed of recording from many thousands of neurons simultaneously. This goal is the major motivation of the BRAIN initiative and related efforts, and with new calcium imaging methods and large - scale multielectrode array (MEA) devices, this dream is quickly becoming a reality. But now a major bottleneck exists. Cutting - edge calcium imaging methods and MEAs output data at rates on the order of *terabytes/hour*, and data rates continue to increase. At these huge rates processing and even storing the data is challenging (Freeman et al 2014 Nature Methods), let alone optimally extracting all the useful information in these data streams; without the right analytical technology, we will never unlock the true potential of these experimental advances.

### Calcium imaging

Calcium imaging has become the dominant method for recording from large populations of neurons, due to several well - known advantages: calcium imaging offers cell - type specificity and can be coupled easily with a variety of genetic tools; imaging approaches can be less invasive and damaging to brain tissue than inserting an MEA; calcium imaging has proven scalability to record simultaneously from O(10^4^) neurons in vivo (over an order of magnitude larger than achieved by an MEA to date); and finally, imaging approaches enable significantly greater experimental design flexibility than MEAs in terms of which subsets of neurons in the imaging volume are interrogated at which times, and how many pixels are assigned to each neuron (we expand on the importance of this point below). At the same time, calcium imaging suffers from some clear disadvantages: calcium signals represent a slow, nonlinear encoding of the underlying spike train signals of interest, and therefore it is necessary to denoise, temporally deconvolve, and spatially demix calcium video data into estimates of neural activity.

There has recently been a flurry of research activity addressing these issues. Building on earlier work (Vogelstein et al 2010; Andilla and Hamprecht 2014), (Pnevmatikakis et al 2016 *; Pachitariu et al 2017; Haeffele and Vidal 2017) present constrained nonnegative or penalized matrix factorization (NMF) approaches to simultaneously solve these demixing and deconvolution problems. (Zhou et al 2016 *) extend this approach to handle data from one - photon imaging approaches, where large “background” contributions from out - of - focus light complicate the demixing problem; recent large - scale approaches to acquiring one - photon imaging data (Bouchard et al 2015; Kim et al 2016) will likely benefit from this approach or modifications thereof. (Friedrich et al 2017a; Giovanucci et al 2017 *) developed real - time implementations that process incoming data online, one imaging frame at a time, enabling closed - loop experiments. (Deneux et al 2016) developed an improved and more general implementation of the hidden Markov model deconvolution approach of (Vogelstein et al 2009). (Picardo et al 2016) developed hierarchical Bayesian methods for sharing statistical information across behaviorally - similar trials to enable temporal super - resolution of estimated neural activity. (Jewell and Witten 2017) developed useful mathematical theory on exactly solving the sparse deconvolution problem addressed in (Pnevmatikakis et al 2016 *; Friedrich et al 2017a). (Wei et al, unpublished, 2017) investigate the impact of calcium indicator nonlinearities on downstream analyses of neural population activity, concluding that some caution is warranted in interpreting neural dynamics inferred from calcium imaging data.

In the near future, we expect that modern computational vision approaches (e.g., based on artificial neural networks (ANNs)) can be incorporated into the NMF framework for further improvements (as we will see below, the incorporation of ANNs to replace modules in various analysis pipelines is a recurring theme in this review); (Apthorpe et al 2016) presented a promising first step. Since NMF is a non - convex problem, accurate initialization of the estimates is critical; the approach presented in (Petersen et al 2017) is a promising alternative approach here (see also Giovanucci et al 2017 *). One major issue that has slowed progress is the lack of “gold standard” datasets that can be used to objectively score algorithm performance. The iterative optimization of open - sourced algorithms on agreed - upon standard datasets has been a critical theme enabling progress in modern machine learning (Donoho 2015 *); see (Berens et al 2017) for a recent application of this general program to improve available calcium deconvolution methods. The curation of “gold standard” spatiotemporal calcium imaging datasets remains a critical challenge; the IARPA MICRONS project (https://www.iarpa.gov/index.php/research-programs/microns) will soon deliver public datasets that combine large - scale electron microscopy with calcium imaging in the same cortical volumes, and will therefore serve as a major step forward in this direction (see also Lee et al 2016; Vishwanathan et al 2017; Bae et al 2017).

One major trend that we see guiding research in this area over the next several years involves the optimization of experimental design and analysis methods jointly in order to image larger populations at higher temporal resolution. It is widely recognized that it does not make sense to e.g. optimize an imaging apparatus in isolation; instead, the full experimental preparation, imaging technology, and computational analysis approach should be considered as parts of a pipeline that should be optimized as a whole. (Yang et al 2016; Prevedel et al 2016; Song et al 2017a; Lu et al 2017 *; Friedrich et al 2017b *) have all offered variations on a similar theme: spatial resolution can be usefully traded off for temporal resolution. That is, we can record from more cells if we are willing to accept a lower ratio of pixels per cell, and, moreover, prior information about cell shapes and locations can shift the favorable point of this trade - off even further: once we know the locations and shapes of the cells in the field of view, we can reduce our spatial resolution even further without negatively impacting the quality of the recovered temporal neural activity (Friedrich et al 2017b *). (Noebauer et al 2017) presented another example of this theme in the context of a computationally - challenging light - field microscopy application; we expect that performance here can be improved significantly with stronger signal models. Simulators such as those developed in (Song et al 2017b) will likely play a useful role in the ongoing joint optimization of demixing methods and hardware design.

One significant problem requires further development: tracking activity with single - neuron resolution in small moving animals with flexible nervous systems, e.g. larval zebrafish (Cong et al 2017), *Drosophila* (Bouchard et al 2015), or hydra (Dupre and Yuste, 2017). Good solutions have been developed in *C*. *elegans* (Nguyen et al 2016, Venkatachalam et al 2016), though demixing of fast cytosolic (non - nuclear - localized) signals remains an unsolved problem. We expect non - rigid registration approaches similar to those developed by (Pnevmatikakis and Giovanucci 2017) to be helpful here; see also (Kim et al 2017 *) for impressive recent progress in larval zebrafish.

While we have focused on calcium imaging in this section, many similar themes will hold for voltage imaging at single - cell resolution, which is expected to be a major growth area over the next decade; see e.g. (Xu et al 2017) for a recent review. Of course voltage imaging also provides the opportunity to record at subcellular resolution, at multiple points along the dendrite and axon; once these imaging methods become more mature we expect to see rapid growth in statistical methods for extracting information from this noisy data, with (Huys et al 2006; Huys and Paninski 2009; Paninski 2010; Paninski et al 2012; Pakman et al 2014) providing a foundation for further development.

### Spike sorting data from large - scale MEAs

Spike sorting has been a not - quite - solved problem for decades. In small - scale recordings, a large degree of manual supervision over the spike sorting process is viable; additionally, it is feasible to manually optimize the depth of a few electrodes or tetrodes to ensure high - SNR recordings. Neither strategy is possible with large - scale MEAs: recordings with hundreds of electrodes are routine now, and much larger MEAs are on the way (Tsai et al 2015; Rios et al 2016; www.darpa.mil/program/neural-engineering-system-design). This looming bottleneck has driven a recent uptick of studies on spike sorting activity from large dense MEAs (Pachitariu et al 2016 *; Yger et al 2016; Jun et al 2017; Lee et al 2017; Dhawale et al 2017; Chung et al 2017); see also (Barnett et al 2016) for an approach for validating spike sorting methods with no ground truth and (Farina et al 2017) for a discussion of similar issues in the context of EMG signals. This recent literature has emphasized computational scalability and proper handling of spike events that overlap across many electrodes. In particular, (Pachitariu et al 2016 *) introduced a fast implementation of a matching - pursuit algorithm to detect these spike overlaps, and (Lee et al 2017) built on this work with a more robust and efficient “triage - then - cluster” approach in which an ANN detects putative spike events and then “clean” spikes are clustered first, followed by more difficult overlapping spikes.

As in the calcium imaging context, it is clear that agreed - upon gold standard datasets will lead to accelerated progress here. Acquisition of ground truth data in this context is a notoriously challenging problem; for now we can only hope for partial gold standard datasets, in which e.g. ground truth spiking for single neurons is available (Neto et al 2016). Optogenetic tagging methods (in which a sparse subset of neurons is activated at known times) could play a very useful role here. In some brain areas we can exploit useful side information to provide a sanity check on the sorting results: for example, the mosaic tiling of receptive fields in the primate retina provides some partial validation. In parallel, as to the imaging context, the iterative improvement of simulators of electrical activity (Hagen et al 2015) remains an important direction. Another useful approach is to create “hybrid” datasets combining simulated spiking signals with real noise signals (Pachitariu et al 2016 *; Jun et al 2017). The time seems ripe for a community - based collaborative approach to develop a battery of gold standard datasets and quality metrics and then iteratively improve each module of these pipelines towards more scalable and accurate solutions.

A separate track of work has taken as a starting point the realization that spike sorting selects the most easily - discriminable units from the observed voltage traces, but leaves behind a large amount of information in the lower - SNR units that can not be separated cleanly from the noise floor. “Clusterless” decoding approaches (Kloosterman et al 2014; Deng et al 2015) have been developed to extract information from these unsorted spikes. A combined strategy (in which we sort the sortable units, but also exploit information from the non - separable units and local field potential signals) works best in practice (Bansal et al 2012; Todorova et al 2014).

Finally, as interest grows in bidirectional electrical neural interfaces that stimulate and record simultaneously, the problem of stimulation artifact cancellation becomes critical; see (Mena et al 2016) for a recent step forward.

## II. Understanding large - scale neural signals

As emphasized above, acquiring and processing large - scale neural data with single - neuron and high temporal resolution has represented a critical bottleneck that has attracted significant research effort over the last couple years. While significant challenges remain, these efforts have established a clear path forward towards eliminating this bottleneck. The next frontier then is to extract understanding from the resulting high - dimensional neural activity data.

Historically, the analysis of spike train data has focused significant effort on three broad questions. (1) How is information encoded in spike trains, and how can we decode this information? (2) Can we infer network connectivity from multi - spike train data? (3) Can we model the activity of large neural populations in terms of a lower - dimensional set of factors? We will review recent progress on each of these three themes in turn below, but it is worth emphasizing up front that models developed to address any of these questions can be profitably combined: e.g., factor analysis models developed to address question (3) can lead to improved decoding of neural data (a la question 1).

### Encoding and decoding

How the brain *encodes* external variables into spike trains, and the converse problem of *decoding* external variables from spike trains, are classic problems in statistical neuroscience. the first problem, generalized linear models (GLMs) have for years provided the methodological foundation: GLMs enable spike trains to be regressed against covariates such as behavioral parameters, hidden factors, and other spiking in the population; see (Paninski et al 2007) for a review. Some recent work has targeted the computational efficiency of GLM estimation methods (Mena and Paninski 2014; Ramirez and Paninski, 2014; Wu et al 2015). Of course, GLMs, being simply a generalization of linear regression methods, have effectiveness dependent entirely on the chosen “feature set” - - that is, the collection of variables against which we choose to regress neural activity. (Chang and Tsao 2017 *) describe an exciting recent application showing that with a good choice of features it is possible to predict highly nonlinear responses. One major trend is to learn features adaptively using a hierarchical approach to combine information over many cells / experiments. This leads to significantly richer and more powerful models. (Field et al 2010) and (Antolik et al 2016 *) are two examples of this approach, in which we share information from simultaneously - recorded cells to estimate a hidden layer that better explains the observed responses. (See also Rahnama Rad et al 2017 for a different method for sharing statistical strength across cells.) Again, modern ANN methods are well - suited to this task of learning a useful shared feature representation from a large dataset of many neural responses: (McIntosh et al 2016, Batty et al 2017 *) provide two examples of this idea (see also Mineault et al 2012 for an earlier example), and we expect to see more applications of this approach in the near future. Alternatively, we can repurpose ANNs trained to perform computer vision tasks (eg object recognition) and use the resulting feature sets to predict responses; see (Kriegeskorte and Diedrichsen 2016?; Yamins and DiCarlo 2016 *) for perspectives on this growing literature.

Regarding the converse problem of decoding, there is a large ongoing engineering and clinical literature on brain - machine interfaces (including not only systems to extract motor control information from the brain but also sensory devices such as cochlear and retinal prosthetics) that we will not attempt to review systematically here. The “ReFIT” decoding algorithm proposed in (Gilja et al 2012) contributed a notable empirical advance in decoding motor signals; (Merel et al 2016) provides a rigorous theoretical foundation and generalization of this algorithm. Another thread of work has shown that stronger models of joint neural variability can be used to construct better decoders (Lawhern et al 2010, Kao et al 2015); see also (Burkhart et al 2016) for a promising converse approach. Finally, ANNs have recently been applied to decoding problems (Sussillo et al 2016; Glaser et al 2017); (Parthasarathy et al 2017 *) notably developed a straightforward procedure for converting an encoding model (i.e., a probabilistic model of the neural responses to an arbitrary stimulus), plus samples from the prior stimulus distribution, into an easily - computed approximation of the optimal Bayesian decoder. We expect to see more applications of similar ideas in the near future.

### Connectivity estimation

Another classic problem in statistical neuroscience is to infer neuronal network connectivity from correlated activity in the network, and then to use the inferred connectivity to understand the For network function and predict its dynamics. The major roadblock here has been the “common input” problem: without strong prior information, it is not possible to reliably distinguish causal connections between pairs of observed neurons versus correlations induced by common input from unobserved neurons. (Soudry et al 2015 *) introduced a novel “shotgun” experimental design that exploits the flexibility of imaging approaches for recording from large populations of cells: the idea is to image different subsets of the network in a serial manner, then use statistical methods to estimate the full network connectivity. (Note that this approach is enabled by optical approaches to neural recording; this approach would not be feasible with current multi - electrode arrays.) In simulations, this approach enables the accurate estimation of networks an order of magnitude larger than was previously possible. (See Turaga et al 2014 for a simplified implementation of this idea.) Experimental methods are now becoming sufficiently fast and scalable to put this method into practice. Other relevant advances include (Harrison et al 2015 *), who introduce new conditional inference methods to address hypotheses about the precision of multineuronal responses, and (Linderman et al 2016), who discuss methods for incorporating stronger prior knowledge into network estimates; see also (Jonas and Koerding 2015 *) for improved prior models for networks.

Once we have estimated the network connectivity, we need a high - throughput method for verifying our estimates (e.g., the inferred synaptic weights). Optogenetic approaches are well - suited to this task; (Shababo et al 2013; Chen et al 2017b) propose a scalable adaptive closed - loop optimal experimental design approach towards mapping and verifying the connectivity onto single postsynaptic cells.

Finally, once these networks are inferred a major goal is to study their dynamical properties. (Gerhard et al 2017; Hocker and Park, 2017) point out that standard GLM estimation approaches can lead to dynamically unstable estimated networks, and propose approaches to correct this deficit; some relevant asymptotic theory is developed in (Hall et al 2017; Chen et al 2017a).

### Factor models

In the language of machine learning, the encoding and decoding problems are *supervised*, in the sense that one seeks a mapping between two known signals: measurable behavioral variables and populations of spike trains. The *unsupervised* analog is often approached using *factor models*: high - dimensional neural population activity is assumed to be a noisy, redundant observation of some hidden (latent), often low - dimensional, signal of interest. This signal can then be interrogated with respect to a scientific hypothesis, used as a denoised and simpler representation of the neural activity, or visualized for compact exploratory analysis of the data.

Following the pioneering work of (Smith and Brown 2003), much of this literature has followed the Bayesian paradigm, where a generative probabilistic model is stipulated to link low - dimensional latent signals to high - dimensional neural spike trains, and then a computational inference procedure recovers the posterior distribution of the latent variable from the observed data. Examples of this paradigm include systems with simple latent temporal structure (e.g., Smith and Brown 2003; Yu et al 2009; Macke et al 2012; Goris et al 2014; Ecker et al 2014; Gao et al 2015; Gao et al 2016 *; Zhao and Park 2016; Zhao and Park 2017), systems with switching dynamical structure (Petreska et al 2012; Kato et al 2015; Wiltschko et al 2015 *; Johnson et al 2016; Linderman et al 2017 *), and systems with recurrent neural network dynamical structure (Krishnan et al 2015; Sussillo et al 2016b). An alternative direct approach to dimensionality reduction is to stipulate an objective or loss function that encodes the features one would like to capture in the data and then optimize a map from data to low - dimensional latent factors (Cunningham and Ghahramani 2015 *).

Dimensionality reduction approaches have been widely used in neuroscience (Cunningham and Yu 2014). Several exemplars of the scientific potential of these approaches are worth noting. First, some of the earliest applications of factor models were to understand mixed selectivity in prefrontal cortex (Machens et al 2010); this work showed that despite the apparent complex responses displayed by single neurons, at the population level simple behavioral correlates can be effectively read out from the brain. Second, one natural but significant extension of this “de - mixing” perspective was the finding that different computations in certain brain areas are carried out in different subspaces of neural population activity, thus providing an implicit gating mechanism for irrelevant activity (Kaufman et al 2014; Elsayed et al 2016 *; Stavisky et al 2017). Third, (Sadtler et al 2014) used this notion of subspaces of activity along with a brain - machine interface to discover constraints (in terms of dimensions in neural population space) on learning. Fourth, population activity along with factor models has been used to understand the dynamical structure of motor and prefrontal cortices (Churchland et al 2012, Mante et al 2013). As more connectomic and cell type constraints become available for population activity recordings, we expect this literature to continue to mature and deepen methodologically, and to elucidate interactions between cell type - specific subpopulations in multiple brain areas (Semedo et. al. 2014; Bittner et. al. 2017).

### Interpretability of Large - Scale Neural Data Analysis

Of course, the impetus behind the analysis of large scale neural data is the belief that these efforts will lead to deeper insights into principles of neural computation. One important line of research, which is in its earliest chapter, is to ask: to what extent is that belief well founded? There are three categories of approach to address this critical question.

First, there is the concern that novel analyses of large - scale neural data may not be discovering new phenomena, but are rather rediscovering simpler, previously known features of the data that appear new given the novel class of data and algorithms used to investigate these points. Recent work has created statistical hypothesis testing frameworks to enable researchers to quantitatively address this rigorous skepticism (Harrison et al 2015 *; Elsayed and Cunningham 2017 *; Laoiza - Ganem et al 2017, Savin and Tkacik 2017), by developing methods to generate datasets that contain these simpler, previously known features but are otherwise random, thus creating a null distribution against which novel large - scale data claims can be tested. Applications to test the presence of linear dynamics in motor cortex (Churchland et al 2012) and the presence of de - mixed readouts in prefrontal cortex (Machens et al 2010) have clarified these previous results (Elsayed and Cunningham 2017 *).

Another key point of skepticism is whether recording larger and larger datasets will produce fundamentally new findings. The answer may depend on the complexity of the experimental paradigm: if the number of recorded neurons grows while the task the animal needs to solve is kept relatively simple, will new scientific insights follow, or must the complexity of the task grow in concordance with that of the data? One group has discussed a notion of required task complexity (Gao and Ganguli 2015), and two others have attempted to measure the complexity of neural population activity in the face of larger and larger datasets, finding both that complexity (as measured by the apparent dimensionality of the data) grows seemingly without bound (Pachitariu et al, unpublished 2017) in some cases, and in others that it does not (Williamson et al 2017). Significant additional theoretical and experimental work is required to provide clearer conclusions here.

Third, at the broadest level, we might ask if our current approaches will ever produce a coherent mechanistic understanding of the neural system. (Jonas and Koerding 2017 *) recently presented a somewhat pessimistic answer to this question; these authors used a man - made computer as a proxy for a small nervous system, then made simulated recordings, applied a battery of statistical analyses, and failed to arrive at a satisfactory understanding of the system’s function or design. Thus their answer to their question, “could a neuroscientist understand a microprocessor,” seems to be negative. While we don’t share the pessimism implicit here, we do agree that despite rapid progress in our field over the last decade, neural data science remains in an early stage, largely because the curve of increasing neural data complexity that we have emphasized above has only recently begun to accelerate sharply upwards. Moreover, many of our theories of the brain have been allowed to flourish largely untethered from data that could constrain and winnow these theories. But now we have to grapple seriously with the question of what we will do when we have in hand, for example, a matrix of the spatially - and temporally - resolved activity of all neurons in an animal performing an interesting behavior. This remains a yet - distant dream in mammals but is close to reality in several invertebrate species, and our field needs to think critically and deeply about what to do now that this century - long goal is almost in our grasp. We believe the way forward is an acceleration of the experiment - analysis - theory cycle; there is a rapidly growing need for new theories to guide our exquisite new experimental tools, and as these theories develop we will continue to need well - matched scalable analysis methods that can connect experiment and theory in a tightly closed loop.

## III. Future outlook

We close by summarizing several trends that will guide development in this field over the next several years.

- Datasets will continue to grow in size as recording modalities are optimized and new approaches are introduced; the scalability of processing pipelines will remain a critical design constraint.
- Closed loop, many - degree - of - freedom optimal control of neuronal populations will represent a critical research subfield as optogenetic spatiotemporal control methods continue to mature (Ahmadian et al 2011; Grosenick et al 2015).
- Fusion of multimodal datasets will represent another critical research area, as large - scale connectomic and cell type constraints (Macosko et al 2015; Lee et al 2014; Kebschull et al 2016) become available to inform functional models (Jonas and Koerding 2015 *).
- With this growth in the scale, quantity, and complexity of datasets and analysis methodologies, statistical techniques for validating and appropriately contextualizing resulting findings will become increasingly essential (Elsayed and Cunningham 2017 *).
- We expect to see more fruitful marriages of “classical” computational neuroscience theories (e.g., network dynamics, reinforcement learning) with statistical models for network inference and dimensionality reduction; see (Linderman and Gershman 2017) for an interesting step in this direction.
- More broadly, sociological trends towards more open and large - scale data sharing and open - source collaborative projects, supported by stronger pipelines (Yatsenko et al 2015) and reproducibility tools (e.g., http://mybinder.org/), will enable richer and more ambitious multilevel models of neural function that are beyond the reach of single laboratories. https://www.internationalbrainlab.com represents an example of this; we expect to see more. Automated curation and compression of data into useful shareable form are important underexplored steps in the analysis pipeline here.
- Finally, from our vantage point the number of critical neural data science projects is currently growing significantly more quickly than the number of young scientists with the necessary interdisciplinary training in machine learning, statistics, and neuroscience. Similarly, as noted above, the richness and complexity of available experimental data is beginning to outstrip the sophistication of the theory that we need to guide new experiments and the development of new analysis approaches. This is becoming a critical bottleneck (Akil et al 2016); increased investment in neural data science and neurotheory training will pay rich dividends in improving our understanding of neural systems over the next decade.

## Acknowledgments

Thanks to Scott Linderman, E.J. Chichilnisky, and Sean Bittner for helpful conversations and suggestions, and Anthony Cruz and Meghan Kase for technical assistance. This work was funded by the DARPA NESD program, NSF BIGDATA IIS - 1546296 (LP), IARPA MICRONS D16PC00003 (LP) and D16PC00008 (LP), NIH NINDS 1U01NS103489 - 01 (LP), NIBIB R01EB22913 (LP), and NEI R21EY02759201? (LP); NIH CRCNS R01 NS100066 - 01 (JPC), the Sloan Research Fellowship (JPC), the McKnight Scholar award (JPC), and the Simons Collaboration on the Global Brain (JPC and LP).

